# Hedgehog dysregulation contributes to tissue-specific inflammaging of resident macrophages

**DOI:** 10.1101/2020.08.16.253237

**Authors:** Mahamat Babagana, Kyu-Seon Oh, Sayantan Chakraborty, Alicja Pacholewska, Mohammad Aqdas, Myong-Hee Sung

## Abstract

Age-associated low-grade sterile inflammation, commonly referred to as inflammaging, is a recognized hallmark of aging, which contributes to many age-related diseases. While tissue-resident macrophages are innate immune cells that secrete many types of inflammatory cytokines in response to various stimuli, it is not clear whether they have a role in driving inflammaging. Here we characterized the transcriptional changes associated with physiological aging in mouse resident macrophage populations across different tissues and sexes. Although the age-related transcriptomic signatures of resident macrophages were strikingly tissue-specific, the differentially expressed genes were collectively enriched for those with important innate immune functions such as antigen presentation, cytokine production, and cell adhesion. The brain-resident microglia had the most wide-ranging age-related alterations, with compromised expression of tissue-specific genes and relatively exaggerated responses to endotoxin stimulation. Despite the tissue-specific patterns of aging transcriptomes, components of the hedgehog (Hh) signaling pathway were decreased in aged macrophages across multiple tissues. *In vivo* suppression of Hh signaling in young animals increased the expression of pro-inflammatory cytokines, while *in vitro* activation of Hh signaling in old macrophages, in turn, suppressed the expression of these inflammatory cytokines. This suggests that hedgehog signaling could be a potential intervention axis for mitigating age-associated inflammation and related diseases. Overall, our data represent a resourceful catalog of tissue-specific and sex-specific transcriptomic changes in resident macrophages of peritoneum, liver, and brain, during physiological aging.

## Introduction

Macrophages are part of the innate immune system and play a major role in antigen presentation, cytokine secretion, and phagocytosis ^1–3^. Tissue-resident macrophages represent self-sustaining and anatomically distinct populations of macrophages that are established early during embryonic development from primitive yolk sac or fetal liver derived cells ^4^. Under normal physiological conditions, resident macrophages maintain tissue homeostasis without contributions from cells of hematopoietic origin. However, notable exceptions include some gastrointestinal macrophages and a subset of peritoneal macrophages, which are derived from hematopoietic progenitors ^4–6^. Tissue homeostasis is compromised with aging due to a low-grade steady state of systemic inflammation, commonly referred to as inflammaging ^7^. Inflammaging, a hallmark of aging, is described as a persistent state associated with elevated levels of pro-inflammatory cytokines, such as TNF-α, IFN-γ, and IL-6 among elderly individuals ^8–11^. This chronic inflammatory condition is thought to increase the risk of many age-related diseases ^12^. While some evidences suggest inflammatory signaling pathways, such as NF-κB signaling, may have increased activity during aging ^13–15^, the signals and mechanisms driving inflammaging remain elusive. Moreover, little is understood about the contribution of tissue-resident macrophages in this progression toward systemic low-grade inflammation.

Here, we investigated age-related characteristics of bone-marrow derived macrophages, peritoneal cavity macrophages, Kupffer cells (liver-resident macrophages), and microglia (brain-resident macrophages), delineating the sex-specific effects of aging on their basal states and responses to endotoxin, using total RNA-seq and secretion profiling of 40 cytokines and chemokines. Our data supports tissue-specific aging among each macrophage population indicative of selective microenvironmental pressures on the aging resident macrophages. Furthermore, we characterized how aging alters the macrophage responses to endotoxin challenge in each biological sex. Our transcriptomic profiling identified reduced hedgehog (Hh) signaling in aged macrophages, which contributed to the basal increase in the expression of pro-inflammatory cytokines, *Tnf* and *Ifng*. These findings suggest that activation of macrophage Hh signaling in the elderly might mitigate chronic age-associated inflammation and promote healthy aging.

## Results

### Age-related changes in the transcriptome of resident macrophages are largely tissue-specific

To investigate the transcriptional changes associated with physiological aging in resident macrophages, we extracted total RNA and performed RNA-seq analysis on basal or lipopolysaccharide (LPS)-treated primary bone marrow-derived macrophages (BMDMs), peritoneal cavity macrophages, Kupffer cells, and microglia from young or aged mice (Figure 1a). We performed a pairwise differential gene expression analysis to compare the two age groups, accounting for biological sex differences (Figure 1b). Two biological replicates were prepared for each experimental group, which confirmed high reproducibility (Figure S1). The largest number of dysregulated genes was observed in microglia with a total of ~200 genes upregulated and ~900 genes downregulated with age. The next largest impact was from Kupffer cells, and peritoneal macrophages and BMDMs had fewer genes with altered gene expression in aging (Figure 1b). Interestingly, age-associated transcriptional changes were largely tissue-specific across the four macrophage populations (Figure 1b). Furthermore, only 7 genes (*Gm16867*, *Hist1h4m*, *1700112E06Rik*, *Gm15446*, *Wdfy1*, *Tmem181b-ps*, and *Dynlt1b*) were commonly dysregulated with aging across all tissue-resident macrophages and in both sexes (Figure S2). This tissue-invariant dysregulation was maintained in response to LPS across all the cell types (Figure S2). These findings indicate that macrophage aging is tissue-specific with few genes dysregulated similarly across these distinct populations.

**Figure 1.**
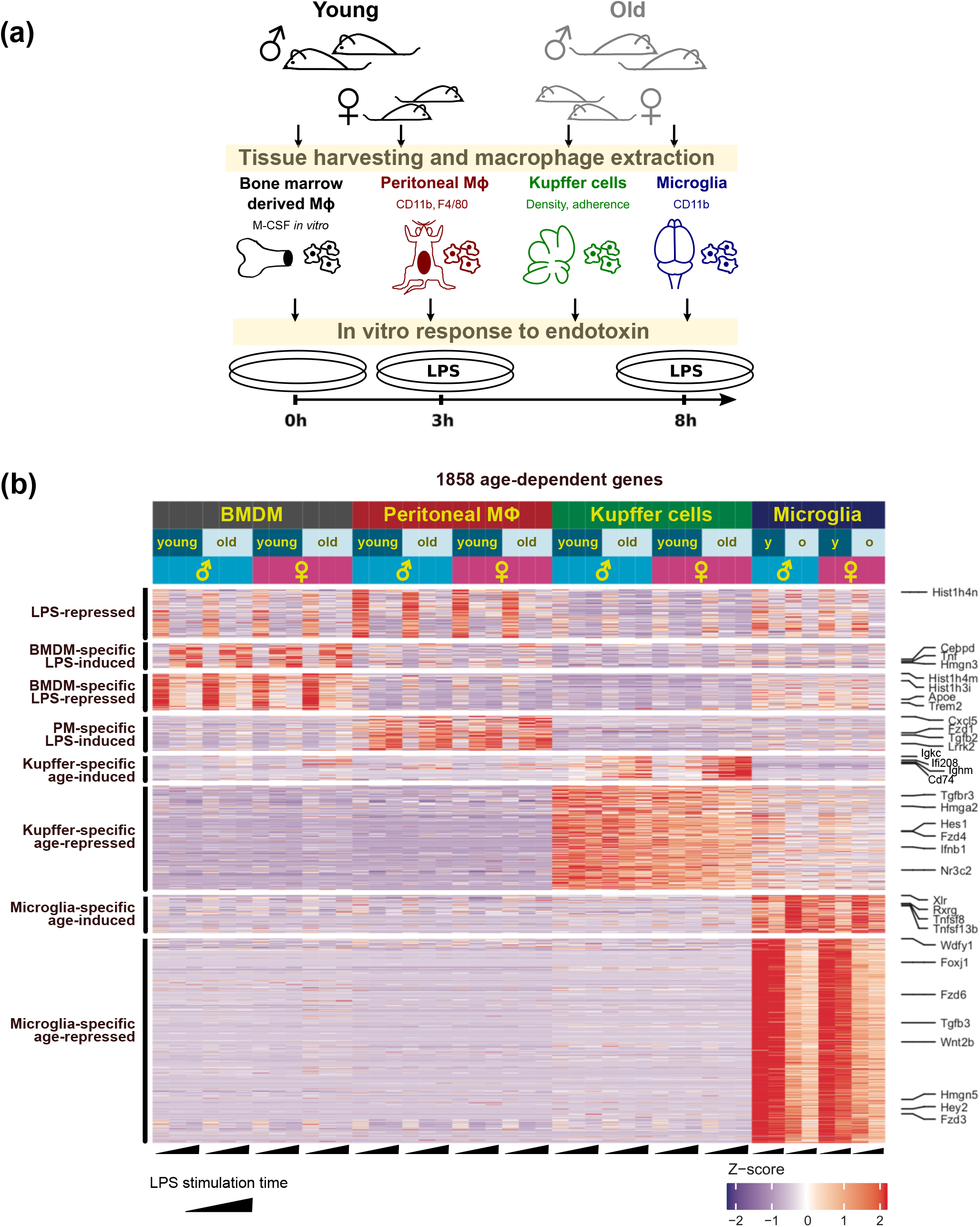
Age-related changes in the transcriptome of resident macrophages are tissue-specific. (a) The workflow with tissue-resident macrophages used to investigate physiological aging in the innate immune system. After processing and treatment, all macrophages were subjected to RNA-seq analysis and other assays as described for each experiment. (b) Clustering analysis reveals tissue-specific gene clusters dysregulated with physiological aging.

### Gene ontology and signature analysis reveals diverse pathways in tissue homeostasis altered in aging

Since dysregulation of the same pathway could be achieved in distinct cell populations through altered expression of different genes, we performed gene ontology analysis to identify any signaling pathways that might be commonly disrupted across resident macrophages in old animals. “Positive regulation of response to external stimulus” and “interleukin-2 production” were the two pathways that were commonly enriched across the macrophage populations (Figure 2a). Furthermore, we observed that the majority of pathways disturbed with aging were related to cell-cell communication and cell-environment interactions, such as cytokine production and cell-substrate adhesion (Figure 2a).This underscores the hallmark of altered cell-cell communication during aging ^8^ and the contribution of tissue-resident macrophages to this phenomenon.

**Figure 2.**
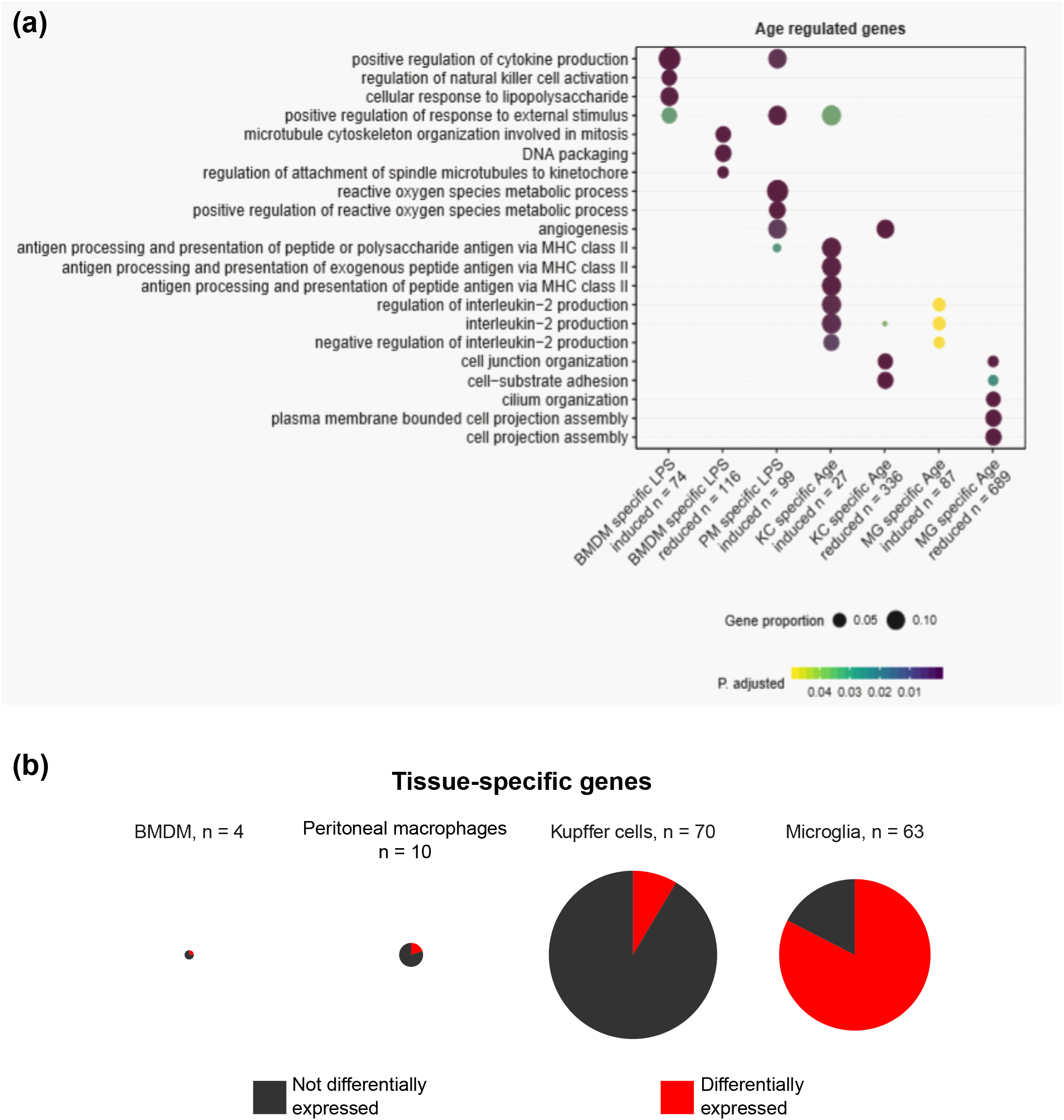
Gene ontology and tissue-specific gene expression signature analysis reveal diverse pathways in tissue homeostasis altered in aging. (a) Gene ontology analysis reveals pathways enriched among dysregulated genes in aging. (b) Pie charts show the proportions of tissue-specific genes unique to each macrophage type that are dysregulated in aging.

Given that age-associated transcriptomic changes are distinctively tissue-specific, we next sought to obtain the distinct tissue signatures of gene expression and asked whether the tissue signatures were maintained during physiological aging. To address this, we identified genes unique to each type of resident macrophages in basal conditions (without LPS). Tissue-specific genes were stringently defined by requiring expression of 10 fragments per kilobase per million reads (FPKM) or higher in the given tissue and lower than 1 FPKM in the other three macrophage types. According to this definition, BMDMs had almost no BMDM-specific genes, and peritoneal macrophages, Kupffer cells, and microglia exhibited 10, 70, and 63 tissue-specific genes, respectively (Figure 2b). In BMDMs, peritoneal macrophages, and Kupffer cells, the majority of tissue-specific genes did not show significant age-associated changes in expression (Figure 2b). However, in microglia, more than two thirds of the microglia-specific genes had altered expression with age. Taken together, the tissue-specific aging of resident macrophages observed in our data supports the notion that the intricate macrophage-niche interactions, rather than an inherent aging program in all macrophages, predominantly shape the aging process. This is also consistent with the fact that distinct tissue environments can drive divergent programs of macrophage gene expression ^16^.

### Tissue-specific impact of macrophage aging drives an altered response to inflammatory stimuli in a sex-dependent manner

To understand how aging affects the macrophage responses to an endotoxin challenge, we analyzed the immediate transcriptomic responses at 3 hours after LPS (10 ng/ml) treatment. Comparison of LPS-to-basal gene expression fold changes between young and old macrophages showed that a large proportion of LPS responsive genes were commonly activated or repressed between the two age groups in each macrophage population. However, a subset of genes was called as differentially expressed in response to LPS treatment in one age group, but not in the other (Figure 3). The transcriptome-wide effects of aging seemed relatively greater in Kupffer cells and microglia, and less in BMDMs and peritoneal macrophages, as assessed by the ‘aging index of gene regulation’: the ratio of the number of age-specific LPS-regulated genes relative to the number of LPS-regulated genes shared between the age groups (Figure 3). Moreover, different genes were age-specifically altered in LPS-responsiveness between males and females. This indicates that sexual dimorphism exists in effects of aging on inflammatory responses, as was observed in age-related metabolic changes and longevity ^17^. For example, our microglia data from male animals showed that *Marco* is an LPS-induced gene that is no longer responsive to LPS in the old. In microglia from female mice, different LPS genes, such as *Csf2*, were disrupted with aging. The aging index of gene regulation for female peritoneal macrophages was nearly twice that of the male counterpart, possibly reflecting the lower replenishment from the bone marrow in females ^18^. To examine the sex differences directly, we obtained sex-specific LPS genes and asked how their regulation changes in aging (Figure S3). In Kupffer cells, the bias of LPS-responsive genes shifted with aging, from male-to female-bias (Figure S3). This analysis showed that LPS-responsive genes had age-dependent patterns of sexual dimorphism. Altogether, these data document a sex-dependent impact of aging on macrophage transcriptomic responses to endotoxin challenge, which manifests distinctively depending on their resident tissue.

**Figure 3.**
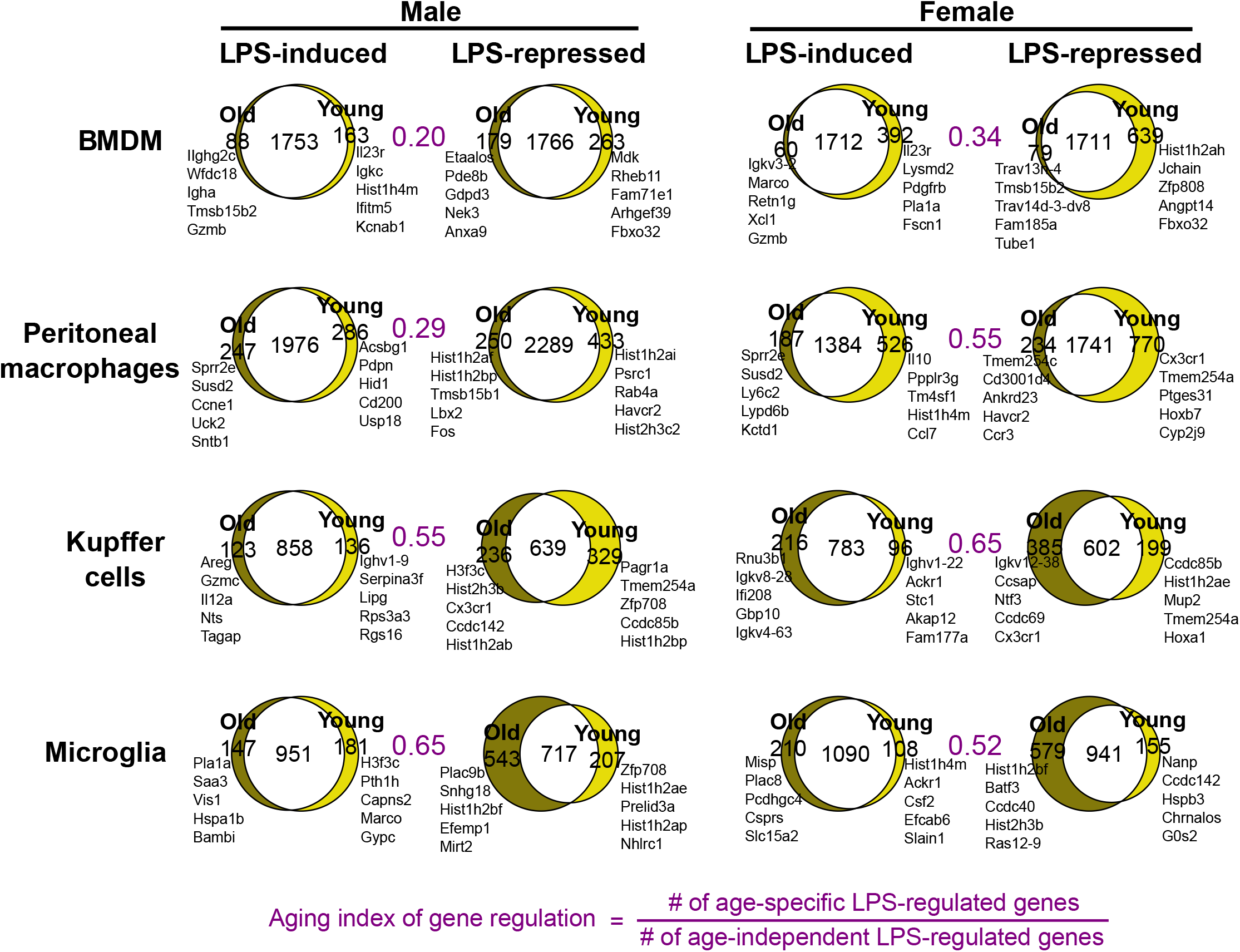
Sex-specific physiological aging generates altered endotoxin responses in tissue-resident macrophages. Venn diagrams show the numbers of LPS-regulated genes common and unique to young and old macrophages. Each set of LPS-induced or -repressed genes was defined as those differentially expressed between macrophages untreated and treated with LPS (10ng/ml) for 3 hours.

### Cytokine profiling reveals tissue-specific secretion patterns in old macrophages

Next, we investigated the functional consequences of the observed transcriptional changes by profiling cytokine secretion. BMDMs, peritoneal macrophages, Kupffer cells, and microglia were plated in the presence or absence of LPS (10 ng/ml) for 8 hours, and supernatant was collected and assayed for the secretion of 40 cytokines and chemokines. Among the resident macrophages, microglia had the most cytokines differentially secreted in old macrophages compared to the young (Figure 4a). Interestingly, the impact of aging was evident in the basal secretion, the induction after LPS, or both conditions, depending on the resident tissue. BMDMs and microglia from old mice showed higher basal levels of distinct cytokines and chemokines, such that the LPS stimulation did not increase the secretion of these cytokines further in BMDMs from old animals. On the other hand, aged microglia exhibited a generally exaggerated response to LPS, showing higher levels of CCL3, CCL4, and CXCL2 in comparison to the old basal or the young LPS-stimulated levels (Figure 4a). Old peritoneal macrophages had a modest basal elevation of a few cytokines including CCL2, but the LPS-stimulated secretion was comparable to the young counterpart. Old Kupffer cells also had a basal increase of CCL2, but the LPS stimulation increased their secretion of many factors, including IL-1ra, CXCL1, CXCL2, CXCL10, CCL2, CCL3, CCL4, CCL5, and TNF-α, well above the levels observed in the young Kupffer cells. Notably, the basal cytokine profile of each resident macrophage population was largely tissue-specific with the exception of TNF-α, CXCL10, and CCL4, which were elevated in all aged macrophages assayed (Figure 4b). These data highlight the selective aging effects on the steady-state cytokine secretion and the endotoxin responsiveness within each tissue.

**Figure 4.**
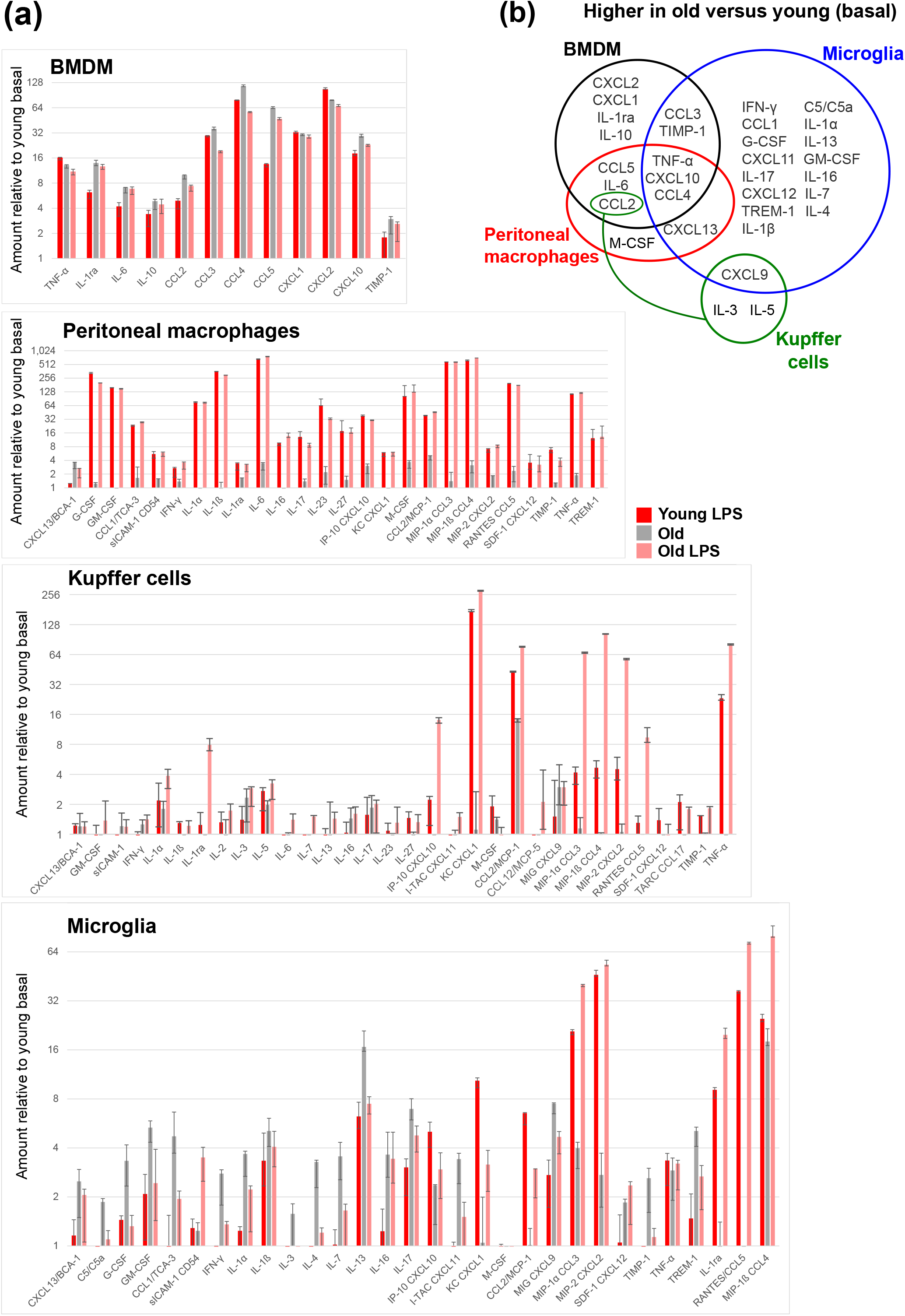
Age-associated changes in cytokine secretion. For each tissue-resident macrophage population in the indicated age group, supernatants were collected from cells untreated or treated with 10 ng/ml LPS for 8 hours, and were subject to a multiplex cytokine detection assay (see Methods). (a) In each plot for the indicated macrophage population, only the cytokines that were detectable in at least one condition are included. (b) A Venn diagram shows the overlap between the cytokines whose secretion was basally elevated with age in each macrophage population.

### Decreased hedgehog signaling in aged tissue-resident macrophages promotes pro-inflammatory cytokine expression

To identify potential pathways contributing to the low-grade basal inflammation in tissue-resident macrophages, we examined genes dysregulated with physiological aging that belong in the same pathways. We noted that the expression of genes involved in Hh signaling were reduced with age across all tissue-resident macrophages. *Smo*, which encodes the key G protein-coupled receptor (GPCR) Smoothened and is involved in the activation of Hh signaling ^19,20^, had significantly reduced expression in old Kupffer cells and microglia (Table S1). Furthermore, *Efcab7*, encoding a protein involved in the localization of the EVC:EVC2 complex and the positive regulation of Hh signaling ^21^, was significantly reduced in peritoneal macrophages and microglia, and a similar trend was observed in BMDMs (Table S1).

Next, we sought to verify whether the decreased expression of Hh signaling components *Smo* and *Efcab7* resulted in a reduction of Hh transcriptional activity. We examined the expression level of canonical Hh target genes *Gli1* and *Ptch1*, which have been shown to positively correlate with Hh signaling ^22^. Although the *Gli1* transcript was undetectable in our dataset, we observed a decrease of *Ptch1* expression in aged peritoneal macrophages, Kupffer cells, and microglia (Table S1). Decreased *Ptch1* expression was also observed in BMDMs, although this did not reach the level of statistical significance (data not shown).

Reduced Hh signaling has been associated with increased expression of the pro-inflammatory cytokine TNF-α ^23^. In agreement with this observation, the basal secretion of TNF-α was increased with aging in BMDMs, peritoneal macrophages, and microglia (Figure 4). Therefore, we asked whether reduced Hh signaling can lead to an increase of pro-inflammatory cytokine production in tissue-resident macrophages. To address this, we inhibited Hh signaling using vismodegib, a potent Smo inhibitor ^24,25^, in young adult mice and assayed for the expression of *Tnf*, the gene encoding TNF-α. A modest but significant increase of *Tnf* was observed in Kupffer cells and microglia extracted from vismodegib-injected animals in comparison to DMSO-injected animals (Figure 5a, b). Furthermore, the expression of *Ifng*, which encodes a cytokine elevated in microglia with aging (Figure 4), was increased several folds in both Kupffer cells and microglia from vismodegib-injected animals in comparison to the control group (Figure 5a, b).

**Figure 5.**
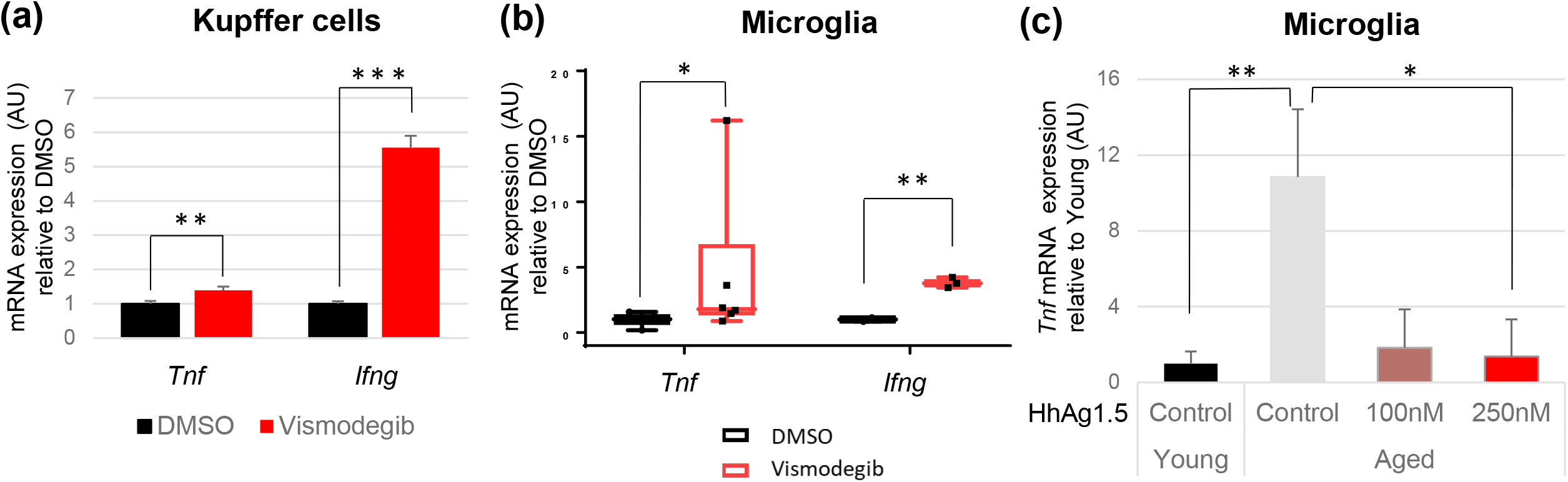
Age-associated reduction of hedgehog signaling in tissue-resident macrophages promotes the expression of pro-inflammatory cytokines. (a) Young mice (2-3 months old) were injected intraperitoneally with vismodegib (75 mg/kg) or DMSO, and RNA was subsequently extracted 1 hour later from Kupffer cells. RT-qPCR was performed to measure the expression of *Tnf* and *Ifng* transcripts. mRNA expression levels were normalized to *Gapdh* and are shown relative to DMSO treated animals. (b) Young mice (2-3 months old) were injected intraperitoneally with vismodegib (75 mg/kg) or DMSO, and RNA was subsequently extracted 6 hours later from microglia. RT-qPCR was performed to measure the expression of *Tnf* and *Ifng* transcripts. mRNA expression levels were normalized to *Gapdh* and are shown relative to DMSO treated animals. (c) Microglia from young or old animals plated for 72 hours were treated with or without HhAg1.5 (Smo agonist) at the indicated doses for 24 hours. *Tnf* expression data measured by RT-qPCR are shown relative to young untreated cells.

Next, we wished to determine if the activation of Hh signaling can reverse the aging-associated low-grade inflammation in aged macrophages. Hence, we isolated microglia from young or aged C57BL/6 mice and treated them *in vitro* with several doses of the Hh agonist, Hh-Ag1.5 ^26^. Aged microglia exhibited the expected increase in *Tnf* expression in comparison to the young. Importantly, Hh-Ag1.5 blocked this increase and the level of *Tnf* expression came down to a baseline comparable to young controls (Figure 5c). Taken together, these results suggest that age-related reduction of Hh signaling in macrophages contributes to basal inflammation, which may be ameliorated when Hh signaling is restored.

## Discussion

Here, we report that the transcriptomic dysregulation associated with physiological aging is largely tissue-specific, which cannot be fully attributed to the altered expression of tissue signature genes in distinct macrophage populations. Only a handful of genes were observed to be commonly dysregulated across all aged macrophages assayed, including those with an uncharacterized function or role in aging. For example, Wdfy1 has been shown to potentiate TLR3 and TLR4 mediated activation of NF-κB and induction of pro-inflammatory cytokines ^27^. However, the lower expression of *Wdfy1* across all the macrophage types in our data may negatively regulate TLR signaling or, alternatively, reflect the state of chronic inflammation (Wdfy1 itself is repressed in response to TLR signaling; Figure S2). Further work is needed to determine the significance of *Wdfy1* dysregulation in the aging phenotype of macrophages across different tissues.

Interestingly, the impact of aging in transcriptomic profiles was most striking in microglia, followed by Kupffer cells. Microglia arise from primitive erythomyeloid progenitor cells during development before mouse embryonic day 8 and are maintained locally in the brain via self-renewal without contributions from postnatal hematopoietic progenitors, unlike some other tissue-resident macrophage populations ^28,29^. Although Kupffer cells and large peritoneal macrophages are established early on during embryonic development, embryonically derived Kupffer cells can be replaced by circulating bone-marrow derived monocytes, and a subset of adult peritoneal macrophages are derived from bone-marrow myeloid precursors ^6,18,29^. We speculate that the magnitude of divergence between the young and old phenotypes may depend on the ability of the tissue-resident macrophages to be replenished by naïve cells from the bone marrow. Nonetheless, even BMDMs, which are *in vitro* differentiated from bone marrow precursors, still showed some aging-related changes in gene expression. This may reflect age-related changes in the niche of the bone marrow which subsequently influences the final cell state.

The majority of pathways dysregulated with physiological aging were related to intercellular communication and environmental homeostasis in our tissue-resident macrophage populations. Interestingly, pathways associated with other notable macrophage functions, such as phagocytosis, were not enriched in our gene ontology analysis of genes that were differentially expressed with aging. It is unclear how aging affects these functions given the varied reports of phagocytosis function among different aged macrophage populations ^30,31^. The profound aging effects in microglia, including the perturbation of two thirds of the genes which are important for their cellular identity, may potentially result from brain-specific changes that manifest more severely, compared to other tissue-resident macrophages. Our data suggest that changes in cell-cell signaling in the microenvironment may be a predominant force that induces age-related alterations of macrophage functions, which is likely to interact with and further influence cell-intrinsic changes. The age-associated changes also manifested in sex-specific patterns, particularly in Kupffer cells and microglia. Notably, the aging index of gene regulation was generally higher in macrophages from females, except for microglia where higher aging index was observed in males.

We demonstrated that aged macrophages have an altered pattern of cytokine secretion in the basal state, which depends on the resident tissue. It is noteworthy that the pro-inflammatory cytokine IL-6, which is detected at higher levels in the plasma of elderly populations ^9–11^, shows increased secretion by some tissue-resident macrophages from our old animals. Elevated levels of TNF-α are also associated with age-related diseases such as dementia and heart disease ^32,33^, and this was indeed one of the few cytokines that were consistently detected to be basally increased across all the aged macrophage populations assayed (Figure 4b). This suggests that these tissue-resident macrophages may be potential sources of increased TNF-α levels during aging ^32,33^. Altogether, our extensive cytokine profiling supports the role of tissue-resident macrophages as major contributors of aged-associated imbalance in pro- and anti-inflammatory factors *in vivo*.

The Hh pathway plays a pivotal role in tissue development, stem cell renewal, and tissue repair ^34,35^. Activation of Smo by mammalian ligands Sonic, Desert, or Indian hedgehog releases the inhibitory effect of Ptch1 and allows Smo to facilitate activation of Gli transcription factors and regulate downstream targets. Aberrant Hh signaling has been observed in several cancers and can act as a driver for tumor angiogenesis, tumor cell proliferation, immunosuppression, and survival ^36,37^. Hh signaling has been reported to stimulate pro-inflammatory cytokine production, while associated with anti-inflammatory properties in other contexts ^23,38–40^. Given the evidence for such dual roles of Hh signaling in controlling inflammation, we had little prior information available to suspect particular involvement of Hh in macrophage aging and their inflammatory phenotype.

Our results support a consistent anti-inflammatory role of macrophage Hh signaling that declines with aging in multiple tissues. To our knowledge, this is the first report that age-associated diminution of Hh signaling increases the expression of pro-inflammatory cytokines. The Hh signaling inhibitor, vismodegib, is currently FDA approved for the treatment of basal cell carcinoma ^41^. Our data support earlier evidence suggesting that the efficacy of Hh inhibition in cancer may be partly due to its inflammatory polarization of tumor-associated macrophages and the infiltration of other immune cells ^23^, in addition to its direct anti-tumor and metastatic properties ^42–44^. The restoration of Hh signaling may potentially delay or prevent age-related diseases associated with chronic inflammation, such as Alzheimer’s disease where resident macrophages play a major role ^45,46^. Interestingly, recent data shows life-extending effect of Hh signaling in AD model of Drosophila^47^. However, given the mitogenic and morphogenic properties of Hh signaling, further research is warranted to investigate whether a balanced and safe Hh signaling regime can be established among the elderly ^34^.

## Experimental Procedures

### Reagents

Collagenase IV (#C2139-1G) and IGEPAL CA-630 (#I8896) were purchased from Sigma. HBSS (#04-315Q) was purchased from Lonza. TGF-beta (#7666-MB) and MCSF (#216-MC-005) was purchased from R&D systems. DMEM/F12 (11320-033), RPMI 1640 (#11835-030), MEM NEAA (#11140-050), HEPES (#15630-080), HBSS with CaCl (#14025-092), Pen Strep (#15070-063), and PBS (#10010-023) were purchased from Gibco. EDTA (#351-027-721) was obtained from Quality Biological. Percoll (#17-0891-01) was purchased from GE Healthcare. LPS (ALX-581-008-L001) was purchased from Enzo. Trizol (#15596018) was purchased from Ambion. Vismodegib (#879085-55-9) was purchased from LC labs. Hh-Ag1.5 (#C4412) was obtained from collagen technology.

### Mice

Young or aged C57BL/6 mice were purchased from Charles River Laboratories. Young mice were 2-3 months of age, and mice at 18-24 months of age were used for the old cohorts. Equal numbers of male and female animals were included, and the different sex samples were processed separately. All mice were maintained under specific pathogen-free conditions at the animal facility of National Institute on Aging, and animal care was conducted in accordance with the guidelines of NIH.

### Tissue and cell extraction

#### Microglia isolation

Whole brains were isolated from adult mice after animal euthanization according to institutional regulations. Tissue dissociation was performed using Miltenyi Biotec’s mouse adult brain dissociation kit (#130-107-677) according to the manufacturer’s protocol. Microglia were purified from total collected cells using CD11b magnetic beads (Mitenyi Biotec #130-049-601) on columns (Miltenyi Biotec #130-042-201). The percentage of microglia among cells was determined by CD45 (Miltenyi Biotec #130-110-632) and CX3CR1 (Biolegend #149039) double-positive cells via flow cytometry. Purity was ascertained to be greater than 90%. Total cells from 3 mice were pooled and used per group (young or aged) and cells were split between the indicated conditions. Equal numbers of male and female mice were used and two biological replicates were performed for each sex. Seeding density was ~0.45E6 cells per 1.9 cm^2^(24-well tissue culture treated dish) with DMEM-F12 in the presence of M-CSF (10 ng/ml) and TGF-β (50 ng/ml). Cells were cultured for 72 hours before subsequent processing. Culture media was supplemented with 10 % FBS and penicillin/streptomycin (1,000 units/ml/1,000 μg/ml).

#### Kupffer cell isolation

Mice were euthanized according to institutional regulations and subsequently transcardially perfused with HBSS for 5 minutes using a peristaltic pump (LKB bromma 2115 multiperpex). The mouse gallbladder was removed before excision of the liver. Livers were subsequently placed and minced in collagenase buffer solution (HBSS, 30 mM HEPES, 1.26 mM CaCl, 0.5 % collagenase IV) for 30 minutes with manual agitation every 15 mins. Cells were collected and centrifuged at 80 g to remove hepatocytes. The remaining cells in suspension were collected and applied to a 50 %/25 % Percoll gradient and centrifuged for ~15 minutes at 800 g with no brake. The interphase was collected and passed through a 70 μm cell strainer. Cells were plated for 1 hour in RPMI supplemented with HEPES (10 mM), 1X MEM Non-Essential Amino Acids, 10 % FBS, penicillin (1,000 units/ml), and streptomycin (1,000 μg/ml). The media was changed after 1 hour to remove any non-adherent cells. Cells were cultured for 1 day before subsequent processing. Cells were seeded at a density of ~0.5E6/1.9 cm^2^(24 well plate). Over 90 % of cells were determined to be F4/80 (Biolegend #123137) and CD11b (Biolegend #101208) double positive by flow cytometry. 2 mice were used for each cohort and 2 biological replicates were performed for each parameter used for RNA-sequencing.

#### Peritoneal cavity-resident macrophage isolation

Peritoneal lavage was performed with 5 ml of freshly-prepared room temperature 0.2 μm sterile-filtered PBS containing 2 mM EDTA and 1 % FBS, per animal. All cells for each sex and age group were collected, pooled, washed, and resuspended in cold DMEM supplemented with 10 % FBS and 2 mM EDTA. Blocking was performed by adding 20 μg/ml anti-CD16/32 (Biolegend #101301) and incubated for 15 minutes on ice in dark. Subsequently, 1.24 μg/ml anti-F4/80 BV421 (Biolegend #123137) and anti-CD11b PE (Biolegend #101208) were added followed by an incubation in dark for 30 min. Cells were washed and resuspended in 500 μl cold DMEM supplemented with 10 % FBS and 2 mM EDTA on ice. ToPro3 (ThermoFisher Scientific #T3605) was added at a dilution of 1:1000 prior to sorting to gate out dead cells. Cells were sorted into 1 ml DMEM supplemented with 20 % FBS. ~0.5E6 cells were subsequently cultured in DMEM supplemented with 10 % FBS in 6-well tissue culture dishes overnight before proceeding to LPS stimulation.

#### Bone marrow derived macrophage

The femur and tibia were removed from young and aged mice. Bone marrow was flushed from each bone using a 27-gauge needle and collected. The cell suspension was passed through a 70 μm cell strainer and collected via centrifugation. Cells were plated in DMEM supplemented with 10 % FBS and M-CSF (60 ng/ml) for 6 days. In the RNA-seq and cytokine secretion experiments, on day 6, BMDMs were plated at 1 million cells per well in a 6-well plate. On the following day, cells were treated with 10 ng/ml LPS and/or harvested as indicated in each experiment.

### RNA extraction and sequencing

Adherent cells were frozen in Trizol and subjected to a freeze thaw cycle prior to RNA isolation. Total RNA was isolated from Trizol/cell lysate mixture by using RNAeasy Mini Kit (Qiagen #74104). Total RNA was sequenced on Illumina NextSeq using Illumina TruSeq Stranded Total RNA library Prep and paired-end sequencing. The stranded paired-end 75 bp reads were mapped to mouse reference genome (GRCm38, mm10 (Church et al., 2011)) with the STAR v2.5.3a mapper (Dobin et al., 2013) using the Gencode M14 primary assembly annotation (Mudge & Harrow, 2015). The reads were counted per gene using RSEM v1.3.0 (Li & Dewey, 2011). Before further analysis, we filtered out the following genes: 1) short genes (< 200bp); 2) ribosomal, transporter RNA genes, and other small RNA genes; 3) genes from unplaced chromosomes. For each cell type, we required a gene to be expressed at 1 FPKM level in at least one sample. Calculated FPKM values were used to test reproducibility among replicates. The count data was then used for differential expression analysis with edgeR v3.18.1 (Robinson, McCarthy, & Smyth, 2010), R v3.4.2 (R Core Team, 2017). Read counts were normalized using the negative binomial generalized linear model distributions for each cell type. Differential expression was determined pairwise for each comparison by likelihood ratio tests using contrasts. A gene was considered differentially expressed if the |log2 fold change| ≥ 0.8 and false discovery rate (FDR) < 0.05. Partition around medoids (PAM) algorithm (Kaufman & Rousseeuw, 1987) was used to draw heatmaps of gene clusters with ComplexHeatmaps R package (Gu, Eils, & Schlesner, 2016). Gene set enrichment analysis was performed using clusterProfiler v. 3.6.0 (Yu, Wang, Han, & He, 2012)

### Western blotting

Whole-cell extracts from BMDM and microglia per condition were prepared in ice-cold RIPA cell lysis buffer (Millipore, 20-188). Cell lysates were centrifuged at 13,000 rpm for 10 min at 4 °C and equal amounts of protein were resolved with a 4 to 20 % Bis-tris Gel/MOPS running buffer system (Invitrogen) and transferred to nitrocellulose membranes. The membranes were analyzed by Western blotting with the following antibodies: rabbit anti-FENS1/WDFY1 (Abcam, ab125329), rabbit anti-Rho-GDI (Sigma, R3025), and horseradish peroxidase (HRP)-conjugated donkey anti-rabbit secondary antibody (GE Healthcare, NA934).

### Drug treatments

C57BL/6 mice of ~2-3 months were intraperitoneally (IP) injected with vismodegib (75 mg/kg) or dimethylsulfoxide (DMSO). Vismodegib was solubilized in DMSO prior to IP injections. Microglia were extracted 6 hours post vismodegib or DMSO injections as described in the cell extraction section of the methods. 6 animals were treated with vismodegib and 5 were treated with DMSO cells to determine *Tnf* expression in microglia. 3 animals were injected with vismodegib and 2 treated with DMSO to determine *Ifng* levels in microglia. Kupffer cells were extracted 1 hour post vismodegib or DMSO injections as described in the cell extraction section of methods. Each animal was treated with vismodegib or DMSO to determine *Tnf* and *Ifng* expression in Kupffer cells.

To determine the effect of HhAg1.5 on macrophages, microglia were extracted and plated as described in the cell extraction of methods. Cells were treated for 24 hours with HhAg1.5 at the indicated doses or untreated. Cells were washed and frozen in trizol. RNA was extracted as described in the RNA extraction section of methods to determine expression of *Tnf* using RT-qPCR.

### Reverse transcriptase-qPCR (RT-qPCR)

cDNA was synthesized from the indicated tissue RNA using iScript cDNA synthesis kit (#170-8891) acquired from Bio-Rad according to the manufacturer’s recommendations. Gene expression was quantified via qPCR using Taqman Probe *Ptch1* (Mm00436026), *Gli1* (Mm00494654_m1), *Hes1* (Mm01342805_m1), *Tnf* (Mm00443258_m1), *Ifng* (Hs00989291_m1) and *Gapdh* (Mm99999915_g1). All gene expression was normalized to *Gapdh* and plotted relative to control.

### Cytokine quantification

Macrophages in male mice were extracted from their indicated tissues and cultured as described above. After the indicated culture period for each type of macrophages, fresh media was added, and the supernatant was collected after 8 hours. Cytokine levels were analyzed using a sandwiched based ELISA array purchased from R&D Systems (ARY006) according to the manufacturer’s protocol. Briefly, 500 μl of supernatant was incubated with a biotinylated-antibody cocktail for 1 hour at room temperature. The antibody/sample mixture was added to a nitrocellulose membrane containing immobilized capture antibodies and incubated overnight at 4 ℃. Streptavidin-HRP and chemiluminescent detection reagents were used to detect each cytokine. Raw pixel density analysis was used to quantity each cytokine level after normalization to each membrane’s reference signal.

### Statistical Analysis

To determine if vismodegib-treated mice significantly increased *Tnf* levels in microglia compared to DMSO treated groups the one-tailed Mann Whitney test was performed; while the Hom-Sidak method was used with GraphPad Prism to assess *Ifng* levels due to fewer biological replicates being performed. *Tnf* and *Ifng* expression in Kupffer cells of mice treated with vismodegib or DMSO was assessed for significance using the student’s t-test in excel. Microglia treated *in vitro* with HhAg1.5 were assessed for *Tnf* expression and compared to untreated cells to determine any statistical difference using the two-tailed Mann Whitney test.

## Acknowledgments

The authors thank Alice Kaganovich and Mark Cookson for their guidance on isolation of microglia from mice. We also thank the National Cancer Institute Center for Cancer Research Sequencing Facility for RNA-seq quality control, library preparation, and sequencing; and the Flow Cytometry core at NIA, Amit Singh, and Arnell Carter for assistance with cell sorting. This work utilized the computational resources of the NIH high performance computing, including the Biowulf cluster. This study was entirely funded by the NIA Intramural Research Program at the National Institutes of Health.

## Author contributions

MB, KSO, SC, AP, MHS designed the study. MB extracted, handled, and processed microglia and Kupffer cells. MB performed and analyzed cytokine assays for BMDMs, peritoneal macrophages, Kupffer cells, and microglia. MB performed the in vivo and ex vivo experiments on microglia. KSO performed the extraction, handling, and processing of BMDMs and performed all western blots. SC extracted, handled, and processed all peritoneal macrophages. AP processed all RNA-seq data, performed subsequent bioinformatics analyses, and generated all the RNA-seq plots. MA performed the Kupffer cell in vivo experiment. MHS supervised all work, provided support and feedback, and completed some data analysis and figures. MB, MHS wrote the manuscript with input from everyone. All the authors edited and approved the final version.

## Competing interests

The authors declare no competing interests.

## Data and materials availability

The materials and data in this study are available upon reasonable request. Raw RNA-seq data were deposited to European Nucleotide Archive (ENA) and are available at https://www.ebi.ac.uk/ena/data/view/PRJEB27853

---

**Figure S1.**
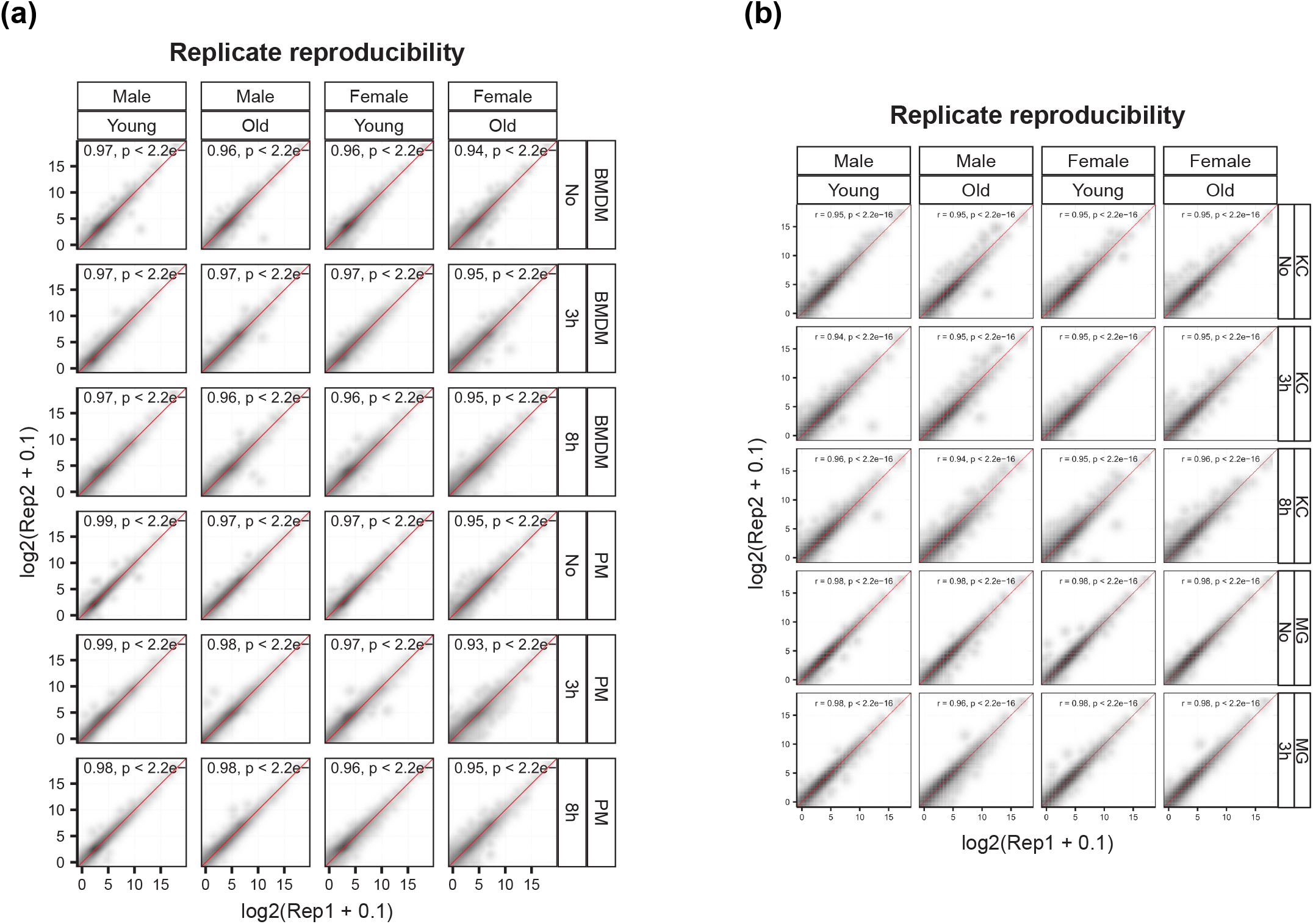
Reproducibility of total RNA-seq between biological replicates. (a) BMDM and peritoneal macrophage (PM) samples. (b) Kupffer cells (KC) and microglia (MG) samples. Rep1: replicate 1, Rep2: replicate 2. Numbers inside each plot are Pearson correlation coefficients and p values.

**Figure S2.**
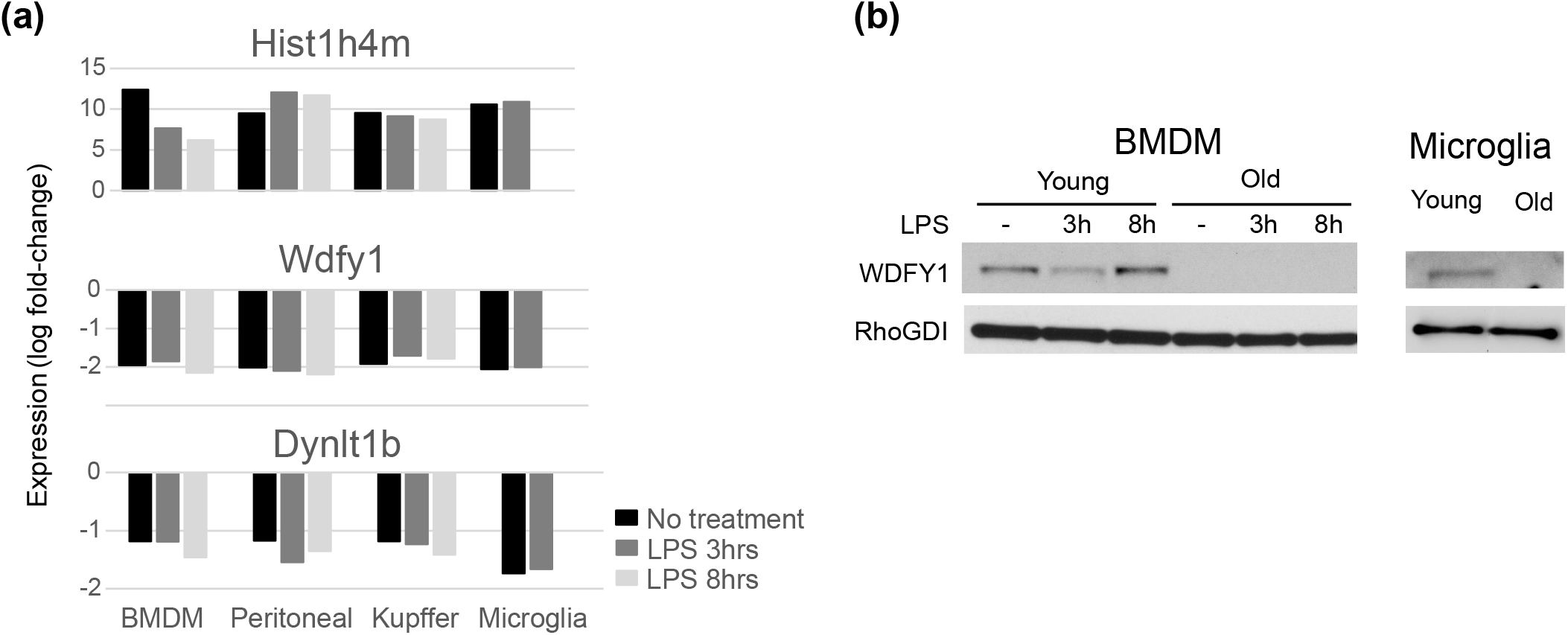
Genes commonly dysregulated with age across the four macrophage populations and all stimulation conditions. (a) RNA-seq expression data are shown in logratios of Old / Young FPKM values. (b) Western blot shows WDFY1 protein expression in BMDM and microglia from young or old mice.

**Figure S3.**
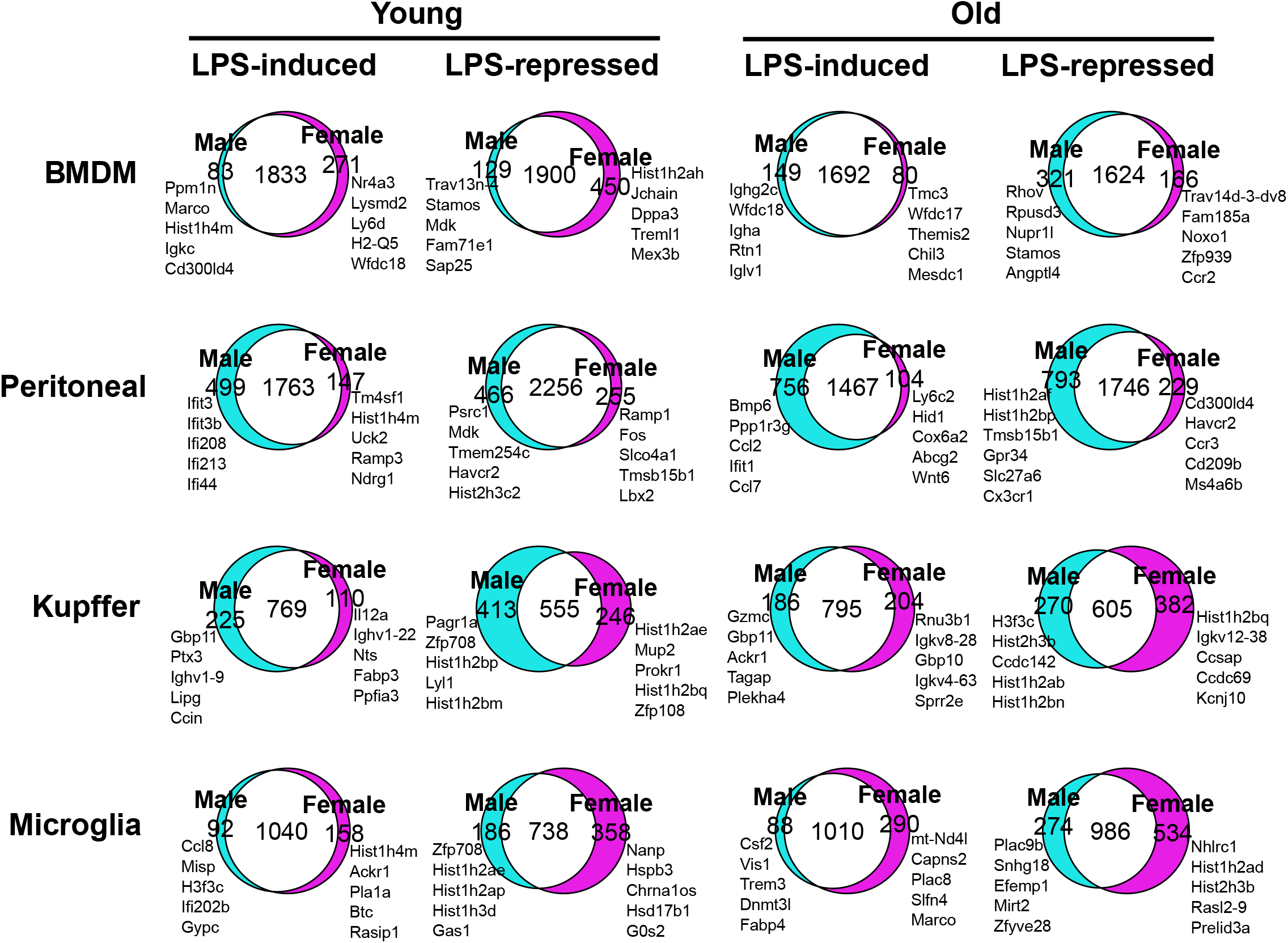
Age-associated sex differences in endotoxin responses of tissue-residence macrophages. Venn diagrams show the numbers of LPS-regulated genes common and unique to male and female mice. Each set of LPS-induced or -repressed genes was defined as those differentially expressed between macrophages untreated and treated with LPS (10 ng/ml) for 3 hours.

## Notes

### Competing Interest Statement

The authors have declared no competing interest.

